# Hippocampal sharp wave ripples mediate generalization and subsequent fear attenuation via closed-loop brain stimulation in rats

**DOI:** 10.1101/2024.04.30.591894

**Authors:** Lizeth Katherine Pedraza, Rodrigo Ordoñez Sierra, Lívia Barcsai, Qun Li, Andrea Pejin, Levente Gellért, Magor Lőrincz, Antal Berenyi

**Author notes:** Corresponding author: Antal Berényi. These authors contributed equally to this work (Lizeth Katherine Pedraza, Rodrigo Ordoñez Sierra).

## Abstract

The balance between stimulus generalization and discrimination is essential in modulating behavioral responses across different contexts. Excessive fear generalization is linked to neuropsychiatric disorders such as generalized anxiety disorder (GAD) and PTSD. While hippocampal sharp wave-ripples (SWRs) and concurrent neocortical oscillations are central to the consolidation of contextual memories, their involvement in non-hippocampal dependent memories remains poorly understood. Here we show that closed-loop disruption of SWRs, after the consolidation of a cued fear conditioning, leads to atypical memory discrimination that would normally be generalized. Furthermore, SWR-triggered closed-loop stimulation of the basolateral amygdala (BLA) during memory reconsolidation inhibits fear generalization and enhances subsequent extinction. Comparable effects were observed when stimulating the infralimbic cortex either post-training or after a brief memory reactivation. A consistent increase in gamma incidence within the amygdala was identified in animals subjected to closed-loop BLA or infralimbic cortex neuromodulation. Our findings highlight the functional role of hippocampal SWRs in modulating the qualitative aspects of amygdala-dependent memories. Targeting the amygdala activity via prefrontal cortex with closed-loop SWR triggered stimulation presents a potential foundation of a non-invasive therapy for GAD and PTSD.

## INTRODUCTION

A delicate balance between an organism’s ability to generalize or discriminate stimuli can prevent inappropriate behavioral responses across contexts. In humans excessive fear generalization has been linked to the development of neuropsychiatric disorders resistant to psycho- and pharmacotherapy such as generalized anxiety disorder (GAD) and posttraumatic stress disorder (PTSD) ^1–3^

Fear conditioning has been used as an animal model to study the most salient aspects of PTSD including fear generalization and the inability to extinguish it ^4–6^. Behavioral interventions with promising results to overcome these issues include extinction, through the formation of a novel memories in competition with the original fear memory ^7^ and reconsolidation, a fundamental plasticity process in the brain that allows established memories to be permanently modified ^8,9^.

In a typical paradigm, conditioned stimuli acquire the property to predict potential dangers, allowing appropriate defense responses. However, overgeneralization to similar but not identical stimuli may result in long-term maladaptive fear expression ^10–12^.

Fear generalization has been associated with neural circuits within the medial prefrontal cortex (mPFC), the anterior cingulate cortex (ACC), the amygdala, and the hippocampus ^11^. In particular, the hippocampus has been shown to play a pivotal role during pattern separation taking place during spatial memory tasks ^13,14^.

Projections from the ACC to the hippocampus are involved in fear generalization of contextual fear memories^15–17^. Indeed, silencing the activity of ACC or ventral hippocampal neurons reduces contextual fear generalization while stimulating these neurons promotes it ^16^. Although the hippocampus is mainly involved in spatial learning ^18,19^, recent studies from our laboratory using closed-loop (CL) stimulation have shown that hippocampal sharp wave ripples (SWRs) are required for the extinction of a cued fear conditioning ^20^, a learning classically linked to the activity of the amygdala ^21–23^. These results are in line with recent reports revealing the hippocampal involvement in long-term memory consolidation previously thought to be hippocampus-independent ^24^. However, as fear extinction is a highly context dependent process itself, the potential role of SWRs in the formation and persistence of cued fear memories remains an open question.

Our results indicate, that closed loop disruption of SWRs through electrical stimulation of the ventral hippocampal commissure (VHC), immediately after the consolidation of a cued fear conditioning, induced the discrimination of memories that would normally be generalized, leading to maladaptive discrimination. In addition, SWR triggered closed-loop basolateral-amygdala stimulation (CL-BLA) prevented fear generalization and enhanced subsequent extinction. The same results were obtained using SWR triggered closed-loop infralimbic cortex (IL) stimulation after training as well as after brief memory reactivation via a reconsolidation-dependent mechanisms. An increase in amygdala gamma oscillation incidence characterized animals stimulated in a closed-loop, but not open loop (OL) regimes. No differences were found in the fear expression to CS+, indicating that only the generalization (response to CS-) was affected.

Overall, our study suggests that hippocampal SWRs participate in the qualitative component of amygdala-dependent memories. This process can be used to update the emotional content of memory traces, reducing generalization, and enhancing subsequent fear attenuation. CL-BLA and closed-loop-medial prefrontal cortex (CL-mPFC) guided by hippocampal activity, emerge as a potential approach to alleviate symptoms of GAD and PTSD.

## RESULTS

### SWRs control memory generalization during the consolidation of cued fear memories

Previous studies from our laboratory have demonstrated that SWRs can be used to update the emotional content of fear memories through closed loop (CL) stimulation of the reward system ^20^. This and other results ^25–27^ highlight the role of SWRs in the processing of fear memories. However, these approaches are performed in tasks where the context acquires a predictive value over the conditioned response (e.g., contextual fear conditioning or extinction). To understand the role of SWRs in tasks that are primarily amygdala-dependent, such as cued fear conditioning^23^ rats underwent a single fear conditioning session, which consisted of five pairings of the conditioned stimulus (CS+) and the unconditioned stimulus (US; 0.8 mA foot shock). Additionally, there were five presentations of CS-without the US (Fig. 1a). After the training, one group of rats was treated with a CL stimulation, characterized by a single 0.5 ms pulse in the VHC. The intensity of this stimulation was set independently for each animal, aiming to disrupt the SWRs and varied between 5–15 V (Fig. 1b). Conversely, OL animals underwent an identical number of randomly timed stimulations within this voltage range, while a control group was not subjected to any stimulation (Fig. 1c).

**Figure 1.**
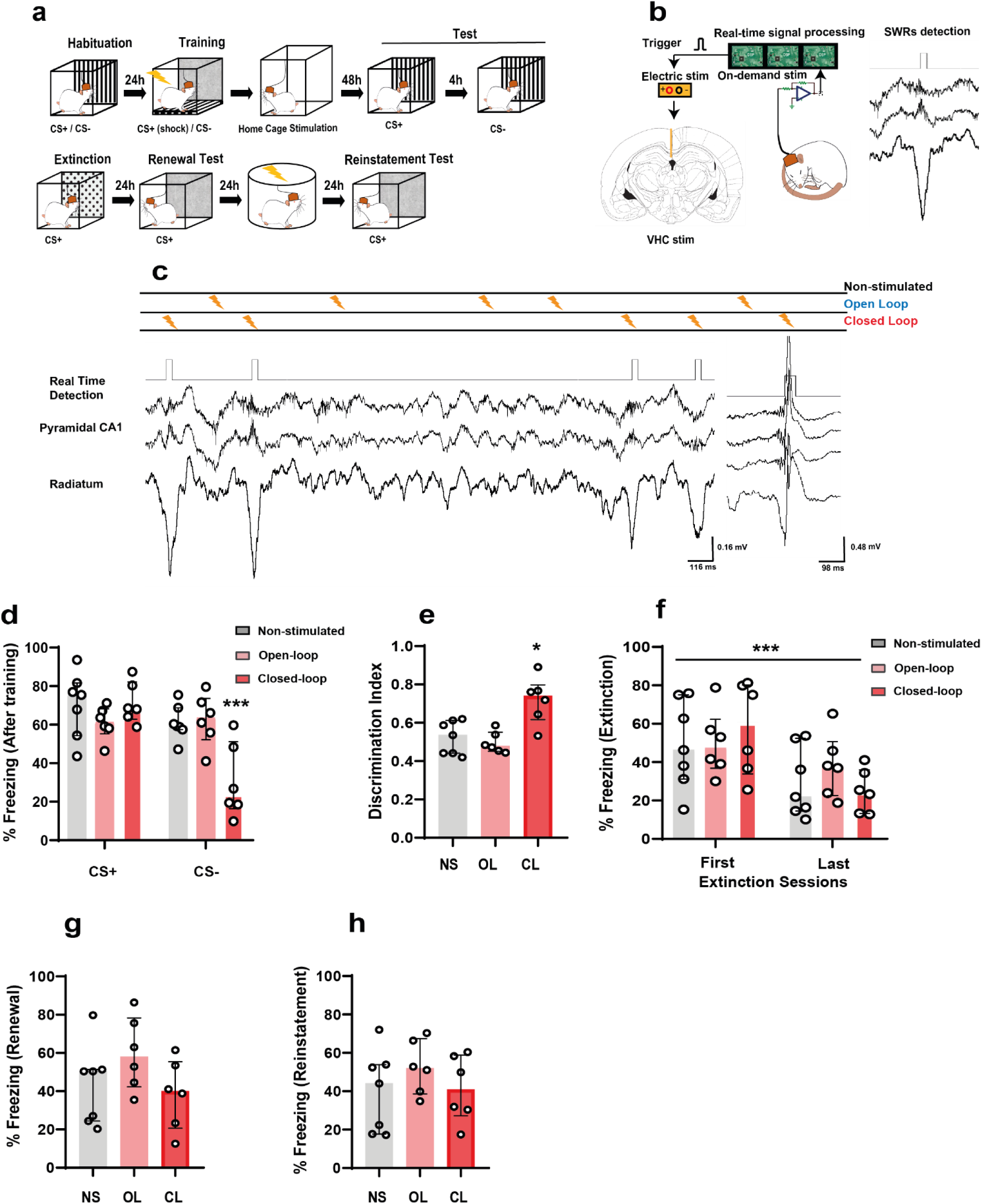
SWRs are required for discrimination of cued fear memories. **a** Schematics of the experimental design. **b** A custom threshold crossing algorithm was used to trigger the VHC stimulation following online detection of SWRs. **c** Closed-loop stimulation (CL) consisted of VHC stimulation during the detected SWR events, open-loop stimulation (OL) was identical to closed-loop but stimulation timing was jittered from SWRs (top). Representative LFP signals from dorsal hippocampus showing intact (left) and disrupted (right) SWRs events. **d** SWR disruption leads to atypical memory discrimination during CS-that would normally be generalized (non-stimulated, NS and open-loop, OL groups) (mixed ANOVA: F (2,16) = 8.739, P < 0.01, groups x time factor (CS + vs CS-) followed by Bonferroni’s multiple comparisons post hoc test (NS vs. CL, P = 0.0009; OL vs. CL, P = 0.0008) (non-stimulated (NS) n = 7; open-loop (OL) n = 6; closed-loop (CL) n = 6). **e** Discrimination indices of the NS, OL and CL conditions (Kruskal-Wallis test: H = 8.415, P = 0.0088). **f** There was a significant decrease in fear expression during extinction (mixed ANOVA: F (1,16) = 20.21, P < 0.0001, time factor), but only CL animas expressed a significant reduction of freezing between the first and last extinction blocks (P < 0.0042). **g** No difference in fear expression during the renewal test (Kruskal-Wallis test: H = 3.155, P = 0.213) and **h** reinstatement test (Kruskal-Wallis test: H = 1.51, P = 0.494). * P < 0.05, *** P < 0.001. Bar plots and error bars represent medians and interquartile ranges, respectively. Individual data points are also displayed. Detailed statistics are shown in Supplementary Data 1. Source data provided as a Source Data file.

The stimulation was performed in the 3 hours immediately following the training. Before initiating the stimulation, a 30 min pre-stimulation baseline was recorded without any interference. To discern the effects of the stimulation, an additional recording of 30 min was conducted post-stimulation. The stimulation was executed within a customized animals’ home cage to reduce the influence of external novelty.

Twenty-four hours later, fear responses to both CS+ and CS-were assessed in separate sessions with an interval of 4 h. On the following day, an extinction session took place, where the rats were exposed to twenty exposures to CS+ in an unfamiliar setting without the US. Post-extinction, the rats were introduced to CS+ test in a hybrid context, combining novel elements with the original conditioning environment (termed the ‘RENEWAL TEST’). This was followed by an unpredictable exposure to the US, and re-test in the hybrid context labeled as ‘REINSTATEMENT TEST’ (Fig. 1a).

In a previous study ^20^, our customized SWR detector demonstrated an 80.38 ± 1.349 % average online success rate compared to the post hoc detection rate. False positive detection rate was 7.750 ± 1.830 %, while the rate of missed detections was 11.88 ± 7.67 %.

Animals subjected to CL stimulation exhibited a reduced freezing level during the CS-test compared to both OL and non-stimulated (NS) groups. Intriguingly, no differences emerged during the CS+ test, suggesting that CL stimulation might specifically affect fear discrimination (Fig. 1d). This observation was further corroborated using a discrimination index, which confirmed that CL-treated animals predominantly restricted fear expression to CS+, whereas the other groups showed generalized fear response (Fig. 1e).

Our results indicate that a single extinction session can induce a general attenuation of fear responses; however, no significant differences were identified between groups (Fig. 1f). Moreover, no differences were encountered between groups during neither the renewal (Fig. 1g) nor the reinstatement phases (Fig. 1h).

In summary, the disruption of SWRs through closed loop neuromodulation during the consolidation phase of cued fear conditioning appears to promote fear discrimination, without affecting the subsequent fear attenuation.

### Closed-loop Basolateral-Amygdala stimulation prevents fear generalization during reconsolidation

The amygdala plays a pivotal role in the processing of fear memories, particularly the consolidation and expression of the cued fear conditioning ^28,29^. Studies in humans suggest a close relationship between amygdala activity and symptomatology in PTSD ^30–32^. Both OL ^33,34^ and CL ^35^ electrical stimulation have proven effective in reducing fear responses in rodents, as well as the severity of symptoms in patients with PTSD. Given the role of hippocampal SWRs in the generalization of cued fear memories, we decided to perform a CL stimulation of BLA and hippocampus. In the subsequent series of experiments, we decided to target the reconsolidation, a second strategy that has been proposed to modify memory traces in a persistent manner ^8,36^. We hypothesized that if SWRs play a dedicate role during the discrimination process, the emotional updating of memory via CL stimulation after reactivation might modify the reconsolidation of cued fear memories.

Forty-eight hours after fear conditioning, which entailed 5 pairings of CS+US at 0.8 mA, animals were introduced to a brief memory reactivation session, involving exposure to 4 CS+, aimed to trigger memory reconsolidation (Fig. 2a). The optimal conditions for this memory reactivation and the verification of reconsolidation were confirmed by a series of parallel experiments (Supplementary Fig. 2). After reactivation, animals underwent a CL-BLA, consisting in a train of 10 pulses, each of 0.2 ms width at 100 μA, 50 Hz, triggered by online detected SWRs (Fig. 2b), animals subjected to OL stimulation experienced the same stimulation, but not cooccurring with SWRs (Fig. 2c).

**Figure 2.**
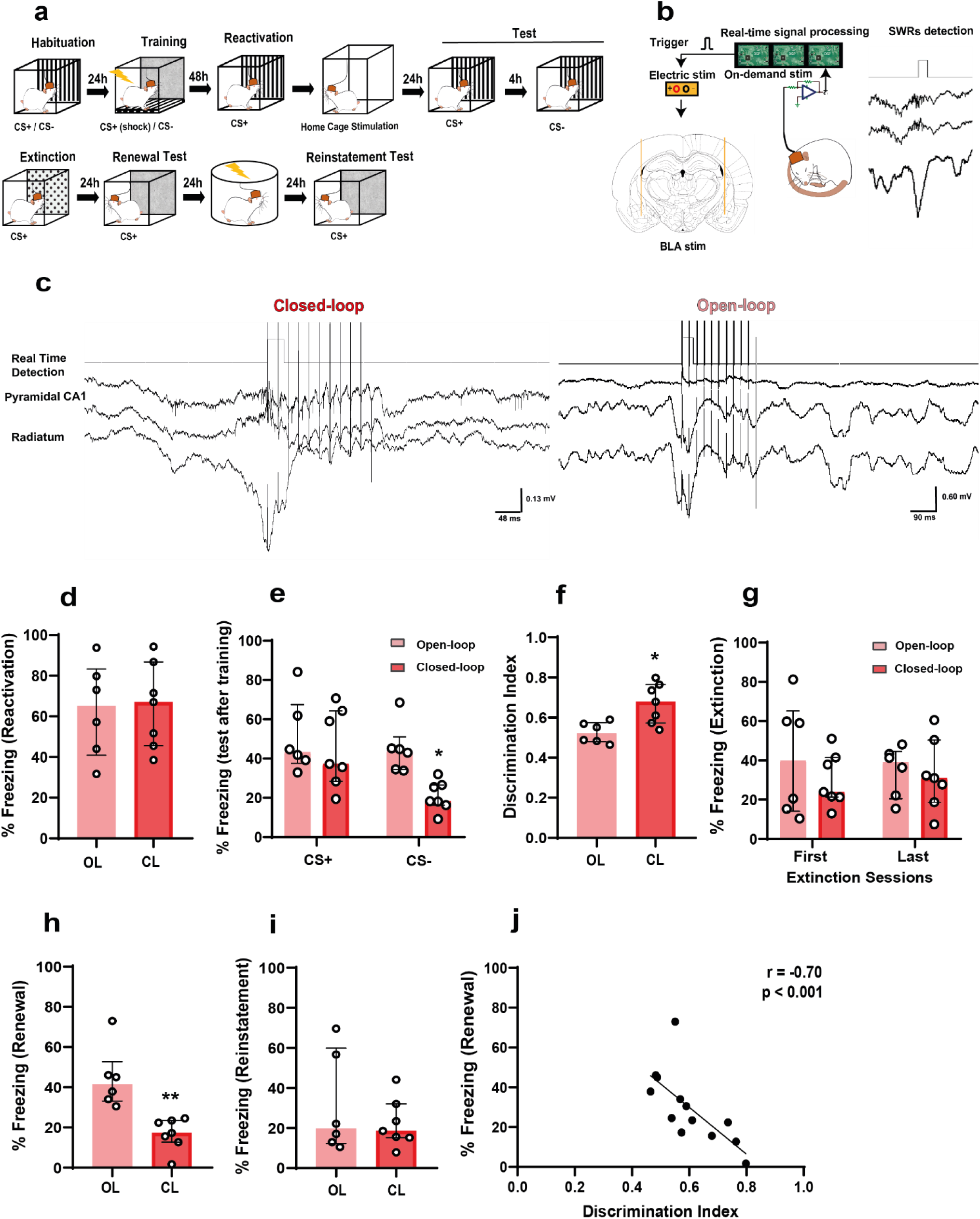
Closed-loop SWR-timed basolateral amygdala electrical stimulation reverts generalization via a reconsolidation-dependent mechanism. **a** Schematics of the experimental design. **b** A custom threshold crossing algorithm was used to trigger BLA stimulation following online detection of SWRs. **c** Representative LFP signals from dorsal hippocampus showing a SWR event and stimulation pattern in closed-loop (left) or random open-loop stimulation (right). **d** No difference in fear expression during the reactivation (Mann Whitney test: U = 20, P = 0.9452, two-tailed; open-loop (OL) n = 6; closed-loop (CL) n = 7). **e** Closed-loop stimulation reverts fear generalization (freezing to CS-) applied after reactivation (mixed ANOVA: F (1,11) = 11.29, P < 0.001, time factor) followed by Bonferroni’s multiple comparisons post hoc test (Test CS-: OL vs. CL, P = 0.0203). **f** Discrimination indices of the NS, OL and CL conditions (Mann Whitney test: U = 4, P = 0.014). **g** No difference was found during the extinction session (mixed ANOVA: F (1,22) = 0.3629, P = 0.5531, group x time interaction). **h** Closed-loop neuromodulation induced lower fear expression during the renewal test in a hybrid context (Mann Whitney test: U = 0, P = 0.0012). **i** No difference was found in the reinstatement test (Mann Whitney test: U = 19, P = 0.8357). **j** Significant negative correlation between the discrimination index and fear expression during renewal (r = −0.70, P = 0.0073). * P < 0.05, ** P < 0.01. Bar plots and error bars represent medians and interquartile ranges, respectively. Individual data points are also displayed. Detailed statistics are shown in Supplementary Data 1. Source data provided as a Source Data file.

During the reactivation phase, no differences were found in fear expression (Fig. 2d). However, similar to the results of Fig. 1, animals that underwent CL-BLA exhibited reduced freezing levels upon exposure to CS-. No differences were identified in responses to CS+ (Fig. 2e), also apparent from the discrimination index (Fig. 2f). No differences were observed during the extinction process (Fig. 2g). Similar to SWR suppression, the closed loop neuromodulation of BLA prevents fear renewal in the hybrid context implying that our intervention may enhance the outcome of fear extinction (Fig. 2h). The reinstatement test showed no differences (Fig. 2i). Interestingly, we identified a significant negative correlation between the discrimination index and fear expression during renewal confirming that preventing memory generalization can enhance fear extinction (Fig. 2j). Discrimination induced by SWRs suppression reflects an impairment in fear memory consolidation, as this correlation was absent during SWRs suppression (Supplementary Fig. 1). Conversely, CL-BLA stimulation reverses fear generalization through a reconsolidation-dependent mechanism and further enhances subsequent fear extinction, potentially through emotional updating ^37–39^. These results are consistent with previous findings about the potential of BLA neuromodulation to achieve fear attenuation ^33–35^.

### Closed-loop IL stimulation reproduce the effects obtained via amygdala stimulation during consolidation

The prefrontal cortex has been shown to be essential for the consolidation and extinction of aversive memories ^40,41^, primarily by exerting an inhibitory effect on the amygdala ^42^. Local inhibition of the IL impairs the extinction and expression of cued fear conditioning memories ^43^, while IL DBS enhances extinction ^44,45^, potentially through decrease in firing rate of principal cells in the BLA ^46^ and inducing anxiolysis ^45,47^. The IL activity, immediately following fear conditioning, seems to control the processes of generalization and subsequent fear extinction ^48^. Collectively, this information leads us to hypothesize that the IL could be a target for less invasive stimulation, while indirectly modulating amygdala activity.

To test this hypothesis, we repeated the memory consolidation experiment of Fig. 1 with IL stimulation (Fig. 3a-b). To control the interaction between memory consolidation and CL stimulation, we implemented a design where a CL stimulation was applied 48 h post-training in an independent group of animals.

**Figure 3.**
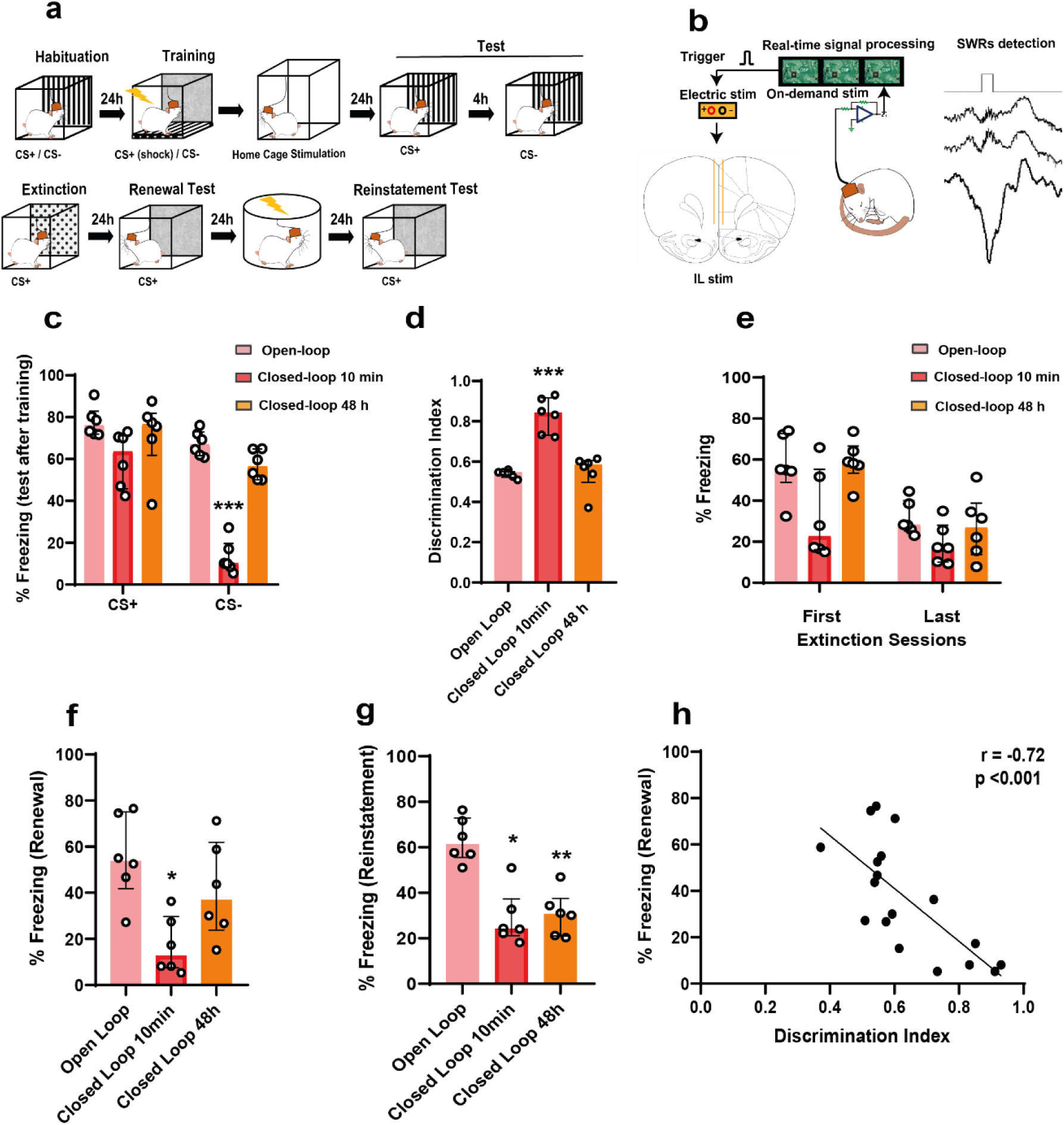
Closed-loop SWR-timed electrical stimulation of the infralimbic cortex prevents generalization applied immediately after training. **a** Schematics of the experimental design. **b** A custom threshold crossing algorithm was used to trigger the IL stimulation following online detection of SWRs. **c** Closed-loop stimulation applied immediately after training, but not 48 h following training reverts fear generalization (freezing to CS-) (mixed ANOVA: F (3,42) = 7.414, P < 0.001, group x tone interaction) followed by Bonferroni’s multiple comparisons post hoc test (NS vs. CL 10 min, P = < 0.001; OL vs. CL 10min, P = < 0.0011; CL 10 min vs. CL 48, P = < 0.001) (non-stimulated (NS) n = 6; closed-loop 10 min (CL-10 min) n = 6; closed-loop 48 h (CL 48 h) n = 6). **d** Discrimination indices of the NS, OL and CL conditions (Kruskal-Wallis test: H = 12.12, P = 0.0002). **e** No difference during the extinction session (mixed ANOVA: F (3,20) = 0.9292, P = 0.4449, group x time interaction). **f** CL stimulation applied immediately following training, but not 48 h later expressed lower fear compared to OL during renewal (Kruskal-Wallis test: H = 8.097, P = 0.0109). **g** Both CL groups expressed lees freezing compared to OL during reinstatement (Kruskal-Wallis test: H = 11.42, P = 0.0005). **h** Significant negative correlation between the discrimination index and fear expression during renewal (r = −0.72, P = 0.0007). * P < 0.05, ** P < 0.01, *** P < 0.001. Bar plots and error bars represent medians and interquartile ranges, respectively. Individual data points are also displayed. Detailed statistics are shown in Supplementary Data 1. Source data provided as a Source Data file.

Similar to the effects of CL-BLA stimulation, animals subjected to immediate post-training CL stimulation displayed reduced freezing reactions to CS-in comparison to both OL and the CL group stimulated 48 h post-training . Responses to CS+ were unaltered (Fig. 3c). This distinction was further confirmed by the generalization index (Fig. 3d). No differences were observed during the extinction session (Fig. 3e). However, in the renewal test, only animals treated with CL immediately post-training showed a significant decrease in fear responses (Fig. 3f). Additionally, during the reinstatement test, both groups (those receiving CL stimulation immediately and 48 h after training) maintained lower levels of freezing compared to the OL group (Fig. 3g). This indicate that CL stimulation itself might enhance the later stages of fear processing. Finally, we identified the same significant negative correlation between the discrimination index and fear expression during renewal as following CL-BLA stimulation (Fig. 3h).

These findings indicates that CL-IL stimulation adequately mirrors the effects observed with BLA stimulation, consistent with previous studies about the modulatory role of IL over BLA ^45,49–51^.

### Closed-loop IL stimulation reverts fear generalization during reconsolidation

While early interventions aiming to update the emotional content or prevent the overconsolidation of aversive memories offered a strategy to reduce the likelihood of PTSD development ^52,53^, neuromodulation systems using electrical stimulation must be readily adaptable to any given moment to ensure feasibility. Memory reconsolidation has been described as a plasticity process that allows previously consolidated memories to be updated ^37,54^, enhanced ^55^, or suppressed ^36,56^. In the case of aversive responses, fear memories suppressed through this reconsolidation method were more resistant to recovery than traditional extinction procedures ^57,58^. Next, we applied the same CL-IL stimulation following a brief memory reactivation capable of inducing reconsolidation processes (Fig. 4a-b). This aimed to overcome the time limitation of applying CL after consolidation and also reduced the invasiveness required for direct modulation of the BLA. Additionally, we introduced a DBS group, which received constant stimulation throughout the 3 h intervention, as previously reported ^45^.

**Figure 4.**
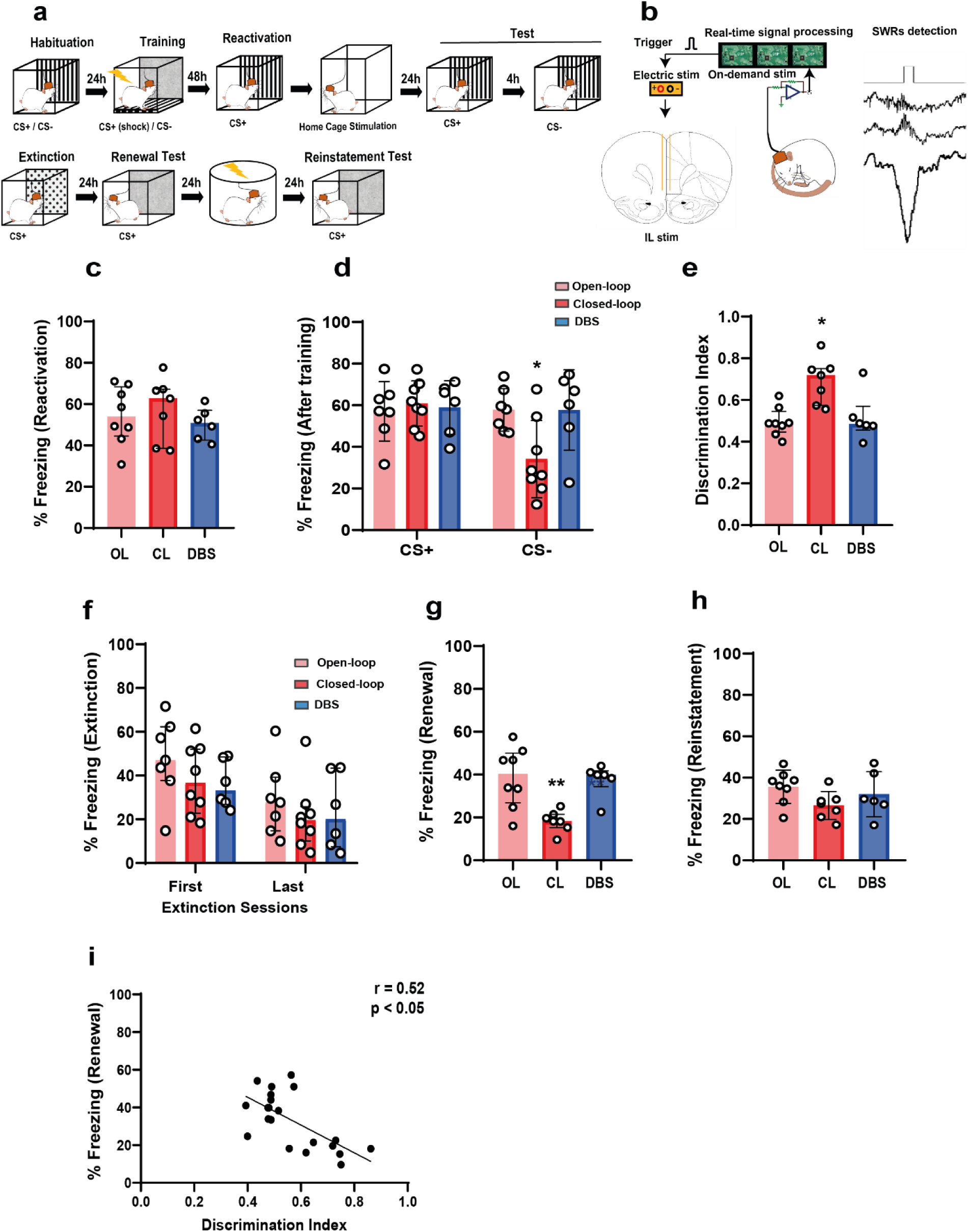
Closed-loop SWR-timed electrical stimulation of the infralimbic cortex reverts generalization via a reconsolidation-dependent mechanism. **a** Schematics of the experimental design. **b** A custom threshold crossing algorithm was used to trigger the IL stimulation following online detections of SWRs. **c** No difference in fear expression during the reactivation (Kruskal-Wallis test: H = 1.068, P = 0.6028) (open-loop (OL) n = 8; closed-loop (CL) n = 7; deep brain stimulation (DBS) n = 6). **d** CL immediately following reactivation, but not 48 h later reverts fear generalization (freezing to CS-) (mixed ANOVA: F (2,18) = 7.905, P < 0.001, group x tone interaction) followed by Bonferroni’s multiple comparisons post hoc test (OL vs. CL, P = < 0.001; CL vs. DBS 10min, P = < 0.01). **e** Discrimination indices of the NS, OL and CL conditions (Kruskal-Wallis test: H = 9.818, P = 0.0034). **f** No difference during the extinction session (mixed ANOVA: F (2,18) = 0.5530, P = 0.5847, group x time interaction). **g** During renewal CL stimulated animals expressed lower fear compared to OL and DBS groups (Kruskal-Wallis test: H = 9.832, P = 0.0033). **h** No difference in the reinstatement test (Kruskal-Wallis test: H = 3.51, P = 0.176). **i** Significant negative correlation between the discrimination index and fear expression during renewal (r = −0.65, P = 0.0012). * P < 0.05, ** P < 0.01. Bar plots and error bars represent medians and interquartile ranges, respectively. Individual data points are also displayed. Detailed statistics are shown in Supplementary Data 1. Source data provided as a Source Data file.

No differences were observed during the reactivation phase in all groups (Fig. 4c). Nevertheless, animals treated with CL stimulation exhibited diminished fear responses when exposed to CS-, compared to both OL and DBS groups (Fig4. d). None of the applied treatments influenced the fear responses to CS+. This enhancement in memory discrimination was also evident in the discrimination index (Fig. 4e). A general decline in fear was noted during the extinction phase, but no significant differences were found between groups (Fig. 4f). Similar to our observations with BLA, CL-IL stimulation during reconsolidation promotes low fear responses during the renewal test (Fig. 4g), with a significant negative correlation between the discrimination index and fear expression during renewal (Fig. 4j). No effects were detected during the reinstatement phase (Fig. 4i).

As observed during consolidation, CL-IL stimulation applied during reconsolidation effectively reproduced the effects of BLA stimulation. By focusing on the prefrontal cortex, our results offer new avenues for non-invasive CL stimulation techniques to accelerate fear attenuation.

### Gamma incidence is enhanced after closed-loop stimulation and correlates with memory discrimination

Several studies suggest that gamma oscillations play a pivotal role in emotional learning and memory ^59–61^. Within the amygdala, particularly in the BLA, gamma rhythms arise from interactions between parvalbumin expressing fast spiking basket cells (PV+) and pyramidal cells ^62,63^. Gamma oscillations have been shown to predict persistent fear memory ^64^. Furthermore, the activation of BLA-PV+ interneurons promotes fear expression and reduces BLA gamma power ^65^. Notably, periods of fear expression correlate with enhanced BLA theta-fast gamma (70–120 Hz) coupling and the suppression of gamma power ^66^, whereas discrimination of safety has been associated with increased BLA fast gamma power ^66^, suggesting a potential substrate for fear discrimination based on gamma rhythms.

Given that both CL-BLA and CL-IL stimulation enhanced fear discrimination, we explored the potential influence of BLA gamma activity on generalization and subsequent fear attenuation. Our results revealed that an increase in gamma incidence post-stimulation is common to both CL-BLA and CL-IL stimulation (Fig. 5b-c), but not observed during VHC stimulation aimed to disrupt SWR occurrence (Fig. 5a). We found a significant positive correlation between gamma incidence and fear discrimination under BLA and IL stimulation (Fig. 5e-f), but not during VHC stimulation (Fig. 5d). Interestingly, conventional DBS seems to decrease gamma incidence (Fig. 5c), yet there is no evidence that this opposite trend leads to impaired discrimination or extinction. In fact, no discernible differences were identified throughout the experiment when compared to the control group (OL) in generalization or fear expression (Fig. 4d-g). These findings suggest that an increase in gamma incidence following CL-BLA and CL-IL might be predictive of fear discrimination and the subsequent enhancement of fear extinction. The lack of this mechanism during VHC stimulation supports our hypothesis that fear discrimination following SWRs suppression originates from a distinct neurobiological substrate, likely associated to the disruption of memory consolidation.

**Figure 5.**
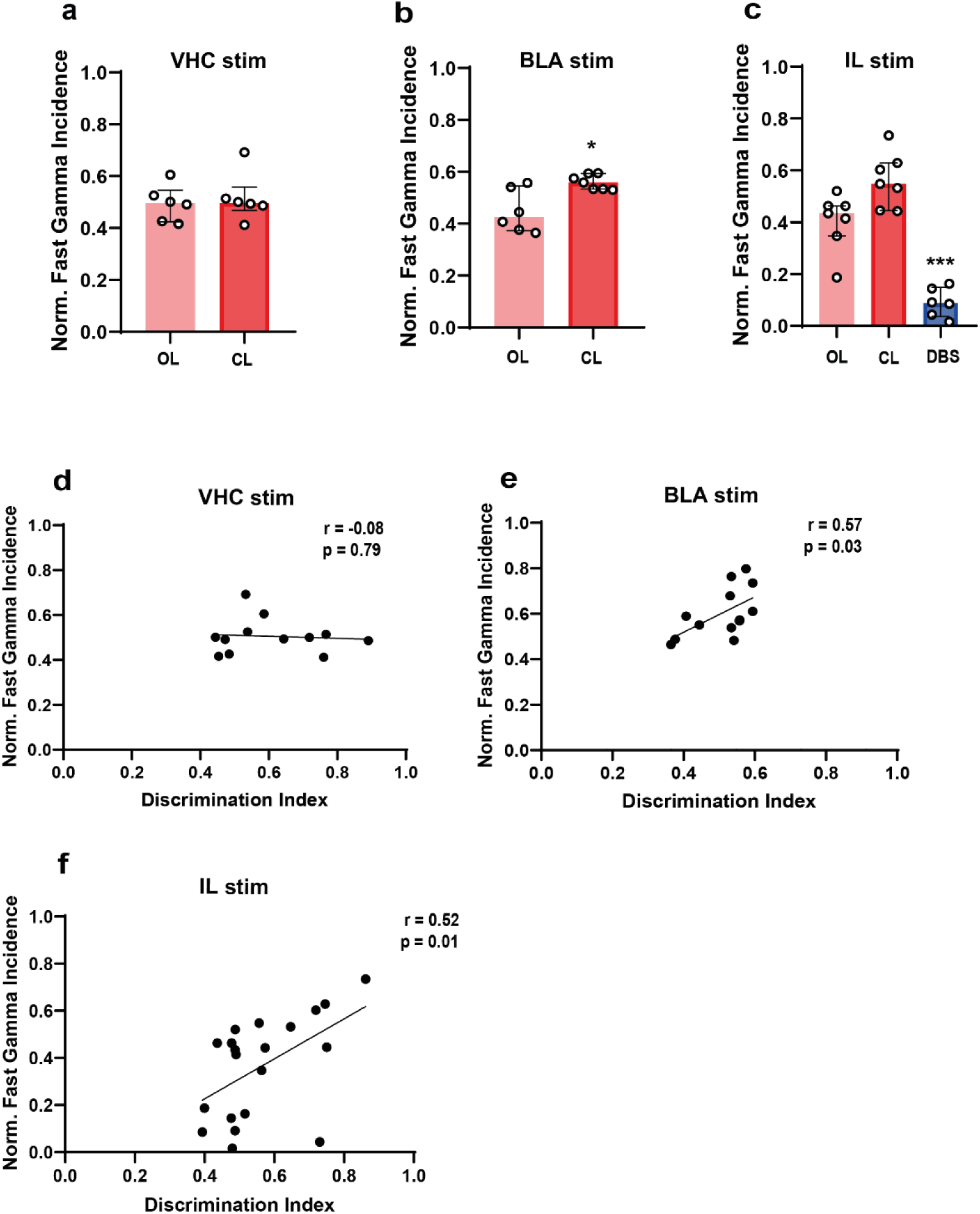
Fear discrimination is correlated with increase of gamma incidence after closed-loop stimulation. **a** No change in BLA gamma incidence following SWRs disruption (Mann Whitney test: U = 18, P > 0.05) (open-loop (OL) n = 6; closed-loop (CL) n = 6). **b** Increased BLA gamma incidence following SWR timed CL BLA stimulation (Mann Whitney test: U = 6, P = 0.035) (open-loop (OL) n = 8; closed-loop (CL) n = 7). **c** Continuous BLA deep brain stimulation drastically reduces gamma incidence compared to closed-loop BLA stimulation (Kruskal-Wallis test: H = 14.79, P < 0.001) (open-loop (OL) n = 6; closed-loop (CL) n = 7; deep brain stimulation (DBS) n = 6). **d** No correlation between gamma incidence after stimulation and the discrimination index in animals submitted to SWR suppression (r = −0.085, P = 0.7916) (Number of values n = 12). However, a significant positive correlation was found CL stimulation of **e** BLA (r = 0.57, P = 0.0377) (Number of values n = 13) and **f** IL (r = 0.52, P = 0.0183) (Number of values n = 20). * P < 0.05, *** P < 0.001. Bar plots and error bars represent medians and interquartile ranges, respectively. Individual data points are also displayed. Detailed statistics are shown in Supplementary Data 1. Source data provided as a Source Data file.

## DISCUSSION

Our study found that SWR suppression can disrupt memory consolidation in amygdala-dependent memories, leading to a non-adaptive memory discrimination without affecting subsequent fear extinction. To our knowledge, this study offers the first evidence that SWRs mediate the precision of non-contextual memories. We revealed that SWRs can serve as a physiological marker to trigger CL-BLA and CL-IL stimulation, thereby reversing the generalization of fear memories through a reconsolidation-dependent mechanism. This approach also prevents the consolidation of generalized fear via IL-stimulation. In both instances, the intervention enhances the subsequent fear extinction process. Notably, a significant negative correlation was found between discrimination and fear expression during renewal under CL-BLA and CL-IL stimulation. These findings are in agreement with an observed increase in gamma incidence following BLA and IL stimulation, suggesting that BLA gamma activity could be a potential predictor for enhanced discrimination and fear attenuation.

Various studies provided strong evidence of the hippocampus and SWRs mediating the consolidation of contextual information, termed as hippocampus-dependent memories ^67,68^ since lesioning or temporally inactivating the hippocampus can affect the learning and retrieval of such information ^69–71^. More recent findings indicate that the hippocampus is also involved in the encoding of non-hippocampal-dependent memories during sleep, as seen in tasks such as novel-object recognition ^24^. Some cortical structures can mediate contextual learning even in the absence of the hippocampus ^72^. Interestingly, a subset of CA1 place cells, respond to specific learning experiences independent of spatial localization ^73^. In line with this, CA2 pyramidal neurons active during a social recognition task can be reactivated during SWRs ^74^.

Here we show that SWRs are crucial in both the consolidation and reconsolidation of amygdala-dependent fear memories, particularly in mediating discrimination. While the disruption of SWRs has been shown to impair the consolidation during spatial tasks ^75^, our results suggest that this manipulation predominantly affected fear expression in response to CS-. Given that none of our CL experiments highlighted differences in response to CS+, we propose that the disruption of SWRs in non-hippocampal-dependent memories affects the qualitative component of the memory, rather than the memory trace itself. We thus propose that the assumed preponderant role of the hippocampus in spatial memories, might be due to tests being biased towards quantifying memory performance based on retrieval success or failure rather than assessing discrimination.

Generalization plays a pivotal role in fear memory processing as a common factor in various anxiety disorders ^76–79^ and a core symptom of PTSD ^80^. Indeed, memory generalization appears to be crucial for prognostic and therapeutic outcomes. For instance, the severity of PTSD symptoms correlates positively with fear generalization ^81^. Moreover, preventing generalization in animal models through chronic antidepressant treatments has shown to enhance subsequent fear attenuation via extinction learning ^82^. It is worth noting that both human and animal studies have shown that females tend to discriminate less compared to males ^83,84^. Additionally, women present a higher risk of developing anxiety-related disorders compared to men ^85,86^.

Interestingly, functional neuroimaging in patients with PTSD has revealed a close relationship between trauma reminders and amygdala activity ^87^. In fact, patients with stress-related disorders exhibit increased amygdala activation in response to fearful faces. This activation correlates positively with a negative recall bias when compared to healthy controls ^88^. A negative bias in memory has been suggested to promote the overgeneralization of negative information ^89^. Intriguingly, animal studies have indicated that amygdala activity might be a predictor for sex differences in fear generalization ^84^. DBS stimulation of the amygdala can mitigate this heightened amygdala activity in animal models of PTSD ^33^, even outperforming traditional antidepressants like paroxetine, which is commonly used to treat PTSD ^34^. These results have been successfully translated to human trials, showing similar outcomes ^90^. It has been proposed that the beneficial effects of DBS amygdala stimulation might be related to the activation of intercalated GABAergic neurons in the BLA, which suppresses the excitatory transmission of principal neurons in key amygdala nuclei responsible for fear expression, such as the central-amygdala ^91^. Consistent with this hypothesis, we propose, that the transient inhibition of the BLA via CL stimulation, aligned with the SWRs, promotes an emotional content update through reconsolidation. The coordinated reactivation between the dorsal hippocampus and BLA during offline memory processing of aversive memories aligns with SWRs ^25^. Considering our findings that SWRs are specifically linked to memory discrimination, SWRs are known to mediate memory processing beyond spatial localization ^73,74^. As a widespread activity characterizes several neocortical and subcortical networks before and during SWRs generation ^92,93^, we suggest that the synchrony of SWRs with precise BLA inhibition via CL results in a less aversive memory and emotional updating, evident in enhanced discrimination. Importantly, we found a significant negative correlation between the ability to discriminate and fear reactions after extinction, suggesting that enhanced discrimination may be a critical step for successful fear attenuation.

Disruption in emotional pattern separation, which relies on interactions between the amygdala and hippocampus, has been proposed as potential consequence of emotional trauma ^94^. Modulating the synchrony between these structures can decrease fear generalization ^95^. Consistent with the role of fast gamma oscillations in the effective discrimination in auditory fear memories ^66^, we observed a notable increase in BLA fast gamma incidence following CL-BLA and CL-IL that correlated positively with the discrimination index. Collectively, these findings indicate that increasing fast gamma activity through CL stimulation can enhance discrimination, which in turn promotes an effective fear attenuation through the extinction process. This phenomenon was not observed during SWRs disruption via VHC CL stimulation.

While BLA CL stimulation timed to SWRs seemed highly effective most likely through the recruitment of transient BLA inhibition, we aimed to explore the possibility that CL stimulation of the infralimbic cortex, a structure amenable for non-invasive stimulation can achieve similar results. The hypofunction of the mPFC leads to maladaptive emotional regulation, including deficits in fear attenuation ^49–51,96^. This hypofunction may result in diminished regulatory activity over the amygdala, aligning with hyper-arousal and heightened fear reactions ^97^. Top-down inhibitory control appears to be compromised in PTSD patients ^98,99^, but can be enhanced through neurofeedback ^100^, psychotherapy^101^, and DBS stimulation ^102^.

In animal models, DBS-IL enhances fear extinction ^44,103,104^. Notably, Reznikov and Colleagues ^45^ demonstrated that chronic DBS-IL ameliorated extinction deficits in animals prone to anxiety-like behavior and subjected to a weak extinction protocol. These findings were mediated by the reduction of BLA neuronal activity in both putative principal cells and interneurons. The role of the IL in fear generalization is less explored, but a recent study suggests that IL activity immediately post-acquisition, but not 6 h later, affects memory generalization ^48^. Furthermore, chemogenetic stimulation of the IL reverses fear memory overgeneralization induced by chronic ethanol treatment in rats ^105^. Given the interaction between the prefrontal cortex and the amygdala during fear reactions, BLA stimulation and CL-IL stimulation could lead to similar behavioral and network effects. This CL stimulation should be performed shortly after learning or reactivation since the same stimulation 48 h post-learning does not enhance fear discrimination.

Collectively, these findings support the idea that transient BLA inhibition during SWRs can be induced either directly through BLA stimulation or indirectly via IL stimulation. The entrainment of fast gamma oscillations in the BLA might be a potential mechanism underlying these outcomes. Whether neurons generating gamma oscillation in the BLA are predominantly recruited during hippocampal SWRs needs to be tested by future studies. Interestingly, when employing a classical DBS stimulation method in the IL, the opposite effect is observed, with a reduction in gamma incidence compared to both CL and OL stimulation. However, no behavioral differences were identified in comparison to the OL group. DBS stimulation might disrupt normal physiological oscillations due to overstimulation and neural habituation ^106^. While DBS targeting both the amygdala and prefrontal cortex has been used to manage PTSD ^90,102^, the adoption of CL versions of these approaches could mitigate potential side effects, for instance epileptiform activity post-BLA stimulation ^91^. Furthermore, CL is based on internal oscillatory feedback activity linked to relevant physiological markers associated with symptoms ^20,35^.

Our findings show significant progress toward the development of entirely non-invasive CL systems to enhance fear attenuation. Given that cortical slow-waves and spindles coincide with SWRs ^107,108^, CL-PFC using tDCS or TMS, triggered by cortical EEG activity, is anticipated to be a viable approach for human trials. Furthermore, future studies should examine these approaches in females, given the extensively reported basal differences in fear generalization compared to males ^84,109–111^.

Overall, our findings suggest that SWRs play a pivotal role in the discrimination of amygdala-dependent fear memories. The temporally precise modulation of both BLA and IL during SWRs can enhance memory precision throughout the consolidation and reconsolidation phases of auditory fear conditioning. Augmenting this emotional pattern separation directly impacts the subsequent fear reduction promoted by extinction. Furthermore, this enhancement is mediated by an increase in BLA gamma incidence. Our methodology offers the scaffolding for feasible non-invasive, CL stimulation treatments for fear and anxiety-associated disorders.

## ACKNOWLEDGMENTS

This work was supported by the Momentum program II of the Hungarian Academy of Sciences (AB), EFOP-3.6.1-16-2016-00008 (AB), EFOP 3.6.6-VEKOP-16-2017-00009 (AB), and KKP133871/KKP20 grants of the National Research, Development and Innovation Office, Hungary (AB), the 20391-3/2018/FEKUSTRAT of the Ministry of Human Capacities, Hungary, the EU Horizon 2020 Research and Innovation Program (No. 739593—HCEMM to AB), Ministry of Innovation and Technology of Hungary grant (TKP2021-EGA-28 to AB), Hungarian Scientific Research Fund (Grants NN125601 and FK123831 to MLL), the Hungarian Brain Research Program (grant KTIA_NAP_13-2-2014-0014 to MLL; and NAP2022-I-7/2022 to AB & MLL), UNKP-20-5 New National Excellence Program of the Ministry for Innovation and Technology from the source of the National Research, Development and Innovation Fund (MLL), Premium Postdoctoral Research Program of the Hungarian Academy of Sciences (RS). MLL was a grantee of the János Bolyai Fellowship.

## AUTHOR CONTRIBUTIONS

L.K.P., R.O.S. and A.B. conceived the project.

L.K.P., R.O.S., L.B., P.A. and G.L. performed the experiments.

L.K.P., R.O.S. and Q.L. analyzed the data.

L.K.P., R.O.S., M.L.L., and A.B. wrote the manuscript.

A.B. supervised the project.

## METHODS

### Animals

60 adult male Wistar, 300-450 g, 3-6 months old were kept in a 12-hour light/ dark cycle. All experiments were performed in accordance with the European Union guidelines (2003/65/CE) and the National Institutes of Health Guidelines for the Care and Use of Animals for Experimental Procedures. The experimental protocols were approved by the Ethical Committee for Animal Research at the Albert Szent-Györgyi Medical and Pharmaceutical Center of the University of Szeged (XIV/218/2016 and XIV/824/2021).

### Surgery

The animals were anesthetized with 2% isoflurane and craniotomies performed according to stereotaxic coordinates. Intracortical electrode triplets (Tungsten 99.95%, California Fine Wire, Grover Beach, CA, USA; interwire spacing, 0.2-0.4 mm) targeting the ACC (AP: +1.0, ML: 0.5, DV: 1.4), bilateral BLA (AP:-2.8, ML: 4.6, DV: 8.1 mm from the dura) and the bilateral CA1 subfield of the dorsal hippocampus (AP: -3.5, -4.5 and -5.5, ML: 2.0, 3.0 and 4.0, DV: 2.9 and 3.0 all mm from the dura). A custom-built bipolar electrode consisting of two insulated (except 200 µm at the tip) Tungsten wires (interwire spacing, 0.2 mm) was implanted in the ventral hippocampal commissure (VHC: -1.3, ML: 0.1, DV: -3.5 mm from the dura), bilateral in the Infralimbic Cortex (IL) (AP: +3.2, ML: 0.5, DV: -4.5 mm from the dura) or bilateral in the BLA (AP:-2.8, ML: 4.6, DV: 8.4 mm from the dura). For combined recording and stimulating in the BLA, stimulus wires were connected together to one pin of connector as anode current input, and five stainless-steel machine screws installed in the skull as cathod and then combined to the other pin of the connector. Stimulation and recording electrodes were secured to the skull with dental acrylic (Unifast Trad, USA). The impedance of the electrodes ranged between 30–90 kΩ at 1 kHz.

Two stainless-steel screws (serving as reference and ground) were placed in the skull above the cerebellum. Some additional screws were drilled into the skull and covered with dental cement to strengthen the implant. A Faraday cage was built using copper mesh where the ground screw was connected and finally, it was attached to the skull around the implanted electrodes with dental acrylic.

In experiments involving pharmacological infusion, rats were bilaterally implanted with 25-gauge guide cannulas, 11mm length (Bilaney Consultants GmbH, Germany) above the BLA (AP: -2.8, ML: 4.7, DV: 6.9 all mm from Bregma). Cannulae were fixed to the skull with dental acrylic (Unifast Trad, USA). The dummy cannula fits 11mm guide without projection to avoid any accidental occlusion.

Post-surgical analgesics and antibiotics were applied lege artis. After recovery, the electrodes were moved in daily steps of 50 to 150 µm until the desired position was reached. Experimental protocol started after a recovery period of at least 15 days following surgery.

### Electrophysiological recordings and stimulation

Rats were housed individually in Plexiglass home cages (42 × 38 cm, 18 cm tall). LFP recordings were conducted in both the home cage and the fear conditioning box (see below). Recording and stimulation sessions for CL or OL interventions were carried out for one or three hours following the training, reactivation, or extinction session, depending on the specific experiment. Food and water were provided ad libitum.

All recording sessions took place in the same room using 12/12 h light/dark cycle with light onset and offset at 7 am and 7 pm, respectively.

The electrodes were linked to a signal multiplexing headstage (HS3_v1.3, Amplipex, Szeged, Hungary) that was connected to a lightweight cable (36AWG Nylon Kerrigan-Lewis Litz wire, Alpha Wire, Elizabeth, NJ, USA. This cable hung from a trolley system on the ceiling of the room, allowing the freely moving. To prevent any twisting or excessive tension in the recording cables, a bore-through electrical commutator (VSR-TC-15-12; Victory-Way Electronic, Shenzhen,China. was used. The multiplexed signals were acquired at 500 Hz per channel for CL neuromodulation experiments26 . The neuronal signals were preamplified (total gain 400X), multiplexed and stored after digitalization at 20-kHz sampling rate per channel (KJE1001, Amplipex, Szeged, Hungary)164. During home cage stimulation, preamplified signals were analyzed on-line by a programmable digital signal processor (RX-8, Tucker-Davis Technologies, Alachua, FL, USA) using a custom made SWRs detection algorithm, as follows.

Two LFP signals were used for real-time SWRs detection. For ripple detection, a channel from the tripolar electrodes from CA1 pyramidal layer with the largest ripple amplitude was selected, band-pass filtered (150–250 Hz), and root-mean square (RMS) power was calculated in real time for ripple detection. For noise detection, manual inspection from channels of the ACC was performed to select the signal with no ripple-like activity and lower noise incidence to enhance signal-to-noise ratio during detection. In case of the ACC, the signal was filtered between 80 and 500 Hz. SWRs were defined as events crossing the ripple thresholds in the absence of the noise signal. Amplitude thresholds for ripple detection were adjusted for each animal before fear conditioning training. Threshold crossings triggered a stimulation (10 pulses, 0.2 ms width at 100 μA, 50 Hz) in the IL and BLA or single pulse in the VHC) (5-15V) (STG4008; Multi Channel Systems, Reutlingen, Germany) depending on the experiment performed. The electrical stimulation of the IL and BLA stimulation was performed under current-controlled mode and VHC stimulation in voltage-controlled mode. Individual thresholds were set for each animal.

### Electrophysiological data analysis

The offline ripples were analyzed using custom-made MATLAB (R2017b, Natick, Massachusetts, USA) routines. Raw signals were down sampled from 20 kHz to 500 Hz and bandpass filtered in the ripple band (150-250 Hz) from the hippocampal channels. Then, normalized squared signals were calculated. Putative SWRs events were defined as those where the beginning/end cutoffs exceeded 2 SDs and the peak power 3 SDs. The detection window was set as 150 ms. SWRs duration limits were set between 20 and 200 ms, otherwise the events were excluded to minimize artifacts. All ripple events were inspected manually following off-line detection. The closest stimulation onset from the digital channel was selected for further analysis. The time delay between the successfully detected ripples events and stimulation time were next quantified. For the brain states classifications (SWS/REM), SleepScoreMaster toolbox from Buzcode (https://github.com/buzsakilab/buzcode) was employed combined with manual corrections. Time frequency spectrum was calculated in MATLAB using Multitaper Spectral Estimation from the Chronux Toolbox (http://chronux.org/). A 2s sliding window with a 50% overlap, a time-bandwidth product of 5 and tapers of 3 were chosen.

Off-line gamma detection was performed for ACC, HPC and BLA. LFP was band-pass filtered with a eighth order zero phase lag Butterworth filter at 30–80 Hz, and RMS power was calculated in 50 ms sliding windows. Outliers of pooled power values were removed to obtain the mean and standard deviation of power values as reference. Gamma bursts were detected where the power values exceeded 3 times of standard deviation (S.D.) above the mean value for gamma frequency for at least three consecutive windows. The boundaries of each gamma events were determined where the power values fell below mean + 2 S.D. around the previously identified peaks. Detection of gamma activity was confirmed by manual inspection. For gamma incidence analysis, awake periods were identified 30 minutes prior to the onset of stimulation and 30 minutes following its conclusion. No stimulation was performed during these times to minimize potential artifact interference. Recording sessions were executed both before and after the three-hour stimulation period, amounting to a total of four hours spent executing the protocol within the home-cage, after either training or reactivation.

### Drugs and infusions

Anisomycin (125 μg/μl; Sigma-Aldrich) dissolved in equimolar HCl and sterile physiological saline (0.9% NaCl) was infused bilaterally into the BLA using a 33G gauge injectors connected to Hamilton syringes via 20-gauge plastic tubes. The infusion injectors tip protruding 2.0 mm below the tip of the cannula and aimed the BLA center. A total volume of 0.5 μl per side was infused by a microinfusion pump at a rate of 0.125 μl/min. Injectors were left in place for an additional minute to ensure proper drug diffusion. The drug was infused immediately after the reactivation session in experiments aimed to confirm that reconsolidation was taking place.

### Auditory fear conditioning

The experiments were performed in a fear conditioning apparatus comprising three contextual Plexiglas boxes (42 × 38 cm, 18 cm tall) placed within a soundproof chamber.4 different contextual configurations were used (Habituation, Reactivation and Test Context (A): square configuration, white walls with black vertical horizontal lines, white smooth floor, washed with 70% ethanol; Training Context (B): square configuration, grey walls, metal grid on black floor, washed 30% ethanol; Extinction Context (C): rectangular configuration, white walls with black dots, white smooth floor; and Renewal and Remote/Reinstatement context (D): hybrid context comprising a square configuration, grey walls from training context, white smooth floor, washed with 70% ethanol. All sessions were controlled using a MATLAB based custom script.

#### Habituation

On day 1, animals were exposed to the habituation session in context A. After 2 min of contextual habituation, they were exposed to 5 alternating presentations of 2 different tones (2.5 or 7.5 kHz, 85 dB, 30 s). Tone time intervals were randomized (30-40 s) during the session. No behavioral differences were detected under exposition to the 2 frequencies.

#### Training

On day 2, cue fear conditioning was performed in context B. After 2 min of contextual habituation, animals received 5 trials of one tone (CS+: 7.5 kHz) immediately followed by a 2 s long footshock as unconditioned stimulus (US: 1.0mA, 0.8 mA or 0.7mA depending on the experiment performed). The other tone (CS−: 2.5 kHz) was presented 5 times intermittently but never followed by the US.

#### Reactivation

Reactivation took place in context A following 2 days after training and entailed four 30 s tone presentations without footshock after an initial 60 s acclimation. Rats remained in the boxes for 60 s after the tone presentation.

#### Test

The tests were performed on days 3-4 and 5-6 in context A. After 2 min of contextual habituation, rats were exposed to presentations of the CS+ or CS-in 2 different sessions. Each session consisted of a block of 5 tones. The order of the CS+ and the CS− in each session was randomized. Sessions were repeated every 4-6 h.

#### Extinction

The extinction was performed on context C, 24 h after the last fear conditioning test. In experiments with manipulations during consolidation and reconsolidation, extinction was performed as a single session consisting of 20 CS+ presentations without the US (unreinforced tones). Tones were repeated with randomized intervals (30-40 s) during the session.

#### Renewal and Remote Test

Animals were exposed to context D (Hybrid context) as a renewal or remote test, respectively, 24 h or 25 days after achieving the remission. In each test, rats were exposed to a block of 5 CS+ presentations after 2 min of contextual habituation. Time intervals between tones were randomized (30-40 s) during the session.

#### Immediate Footshock

To promote fear recovery, animals were placed in a neutral environment outside the conditioning box and received an unconditioned foot shock after 30 s contextual exposition, with the same intensity used during fear conditioning. The animals were returned to their home cage 30 s following the footshock.

#### Reinstatement Test

Animals were submitted to a reinstatement test in context D 24 h after the immediate footshock. Rats were exposed to a block of 5 CS+ presentations without the US after 2 min of contextual habituation. Time intervals between tones were randomized during the session.

### Behavioral Assessment

The measure of freezing behavior was used as an indicator of memory in the fear conditioning task. Freezing was analyzed off-line using Solomon software (SOLOMON CODER, (© András Péter, Budapest, Hungary), for behavioral coding by an experienced observer that was blind to the experimental group. Freezing was defined as the absence of all movements, except those related to breathing, while the animal was alert and awake. An index discrimination analysis between tones was used based on previous studies: [Freezing CS+/(Freezing (CS+) + Freezing (CS-))] (Pedraza et al., 2017).

### Histology

Following the termination of the experiments, animals were deeply anesthetized with 1.5 g/kg urethane (i.p.) and the recording sites of each electrode were lesioned with 100 µA anodal direct current for 10 s. Then, the animals were transcardially perfused with 0.9% saline solution followed by 4% paraformaldehyde solution and 0.2% picric acid in 0.1 M phosphate buffer saline. After postfixation overnight, 50-μm thick coronal sections were prepared with a microtome (VT1000S, Leica), stained with 1 µg/ml DAPI in distilled water (D8417; Sigma-Aldrich), coverslipped and examined using a Zeiss LSM880 scanning confocal microscope (Carl Zeiss) for histological verification of the recording electrode and cannulae locations (Supplementary Fig. 3).

### Statistical analysis and data presentation

Statistical analyses were performed using GraphPad Prism 8 software. Significance was set at p < 0.05. Data were analyzed using two-tailed Mann–Whitney U test, Kruskal–Wallis test or ANOVA Mixed-Effect Analysis followed by Dunn’s post hoc or Bonferroni’s multiple comparisons test. Data are expressed and visualized as median ± IQR, individual data points are also shown where applicable. *, ** and *** denote significance levels smaller than 0.05, 0.01 and 0.001, respectively.

## DATA AVAILABILITY

The data generated in this study (in the main manuscript and in the Supplementary Information) are provided in the Source Data file and Supplementary Data 1, or from the corresponding author upon reasonable request. Source data are provided with this paper.

## CODE AVAILABILITY

All custom code is freely available from the corresponding author on reasonable request.

