## Supplementary Fig for "Hippocampal sharp wave ripples mediate generalization and subsequent fear attenuation via closed-loop brain stimulation in rats"

*[1]MTA-SZTE 'Momentum' Oscillatory Neuronal Networks Research Group, Department of Physiology, University of Szeged, Szeged, 6720, Hungary; [2]HCEMM-SZTE Magnetotherapeutics Research Group, University of Szeged; Szeged, 6720, Hungary; [3]Neunos ZRt; Szeged, 6720, Hungary; [4]Department of Physiology, University of Szeged, Szeged, 6720, Hungary; [5]Neuroscience Division, Cardiff University, Museum Avenue, Cardiff CF10 3AX, UK; [6]Neuroscience Institute, New York University; New York, NY 10016, USA*

† These authors contributed equally to this work

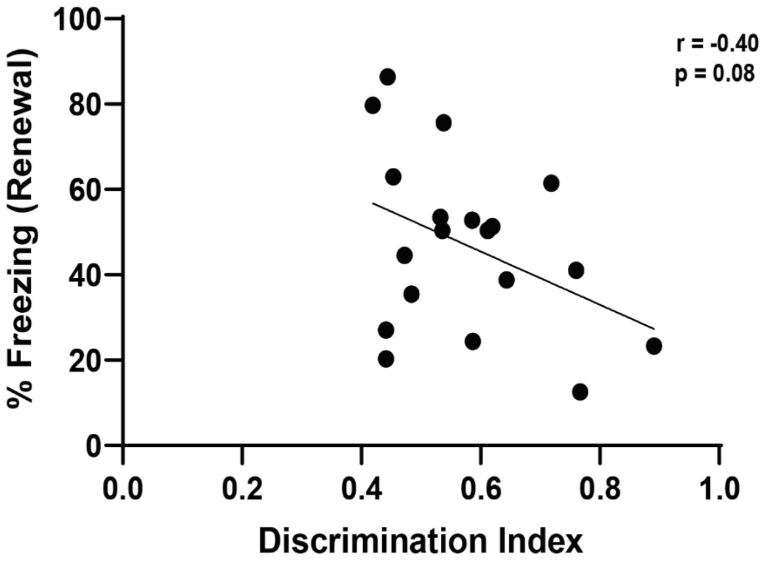


**Supplementary Fig. 1. Enhanced discrimination after SWR disruption is not correlated with fear expression during the renewal.** No significant correlation between the discrimination index and fear expression was detected during renewal following SWR disruption (r = −0.40, P = 0.08).

a

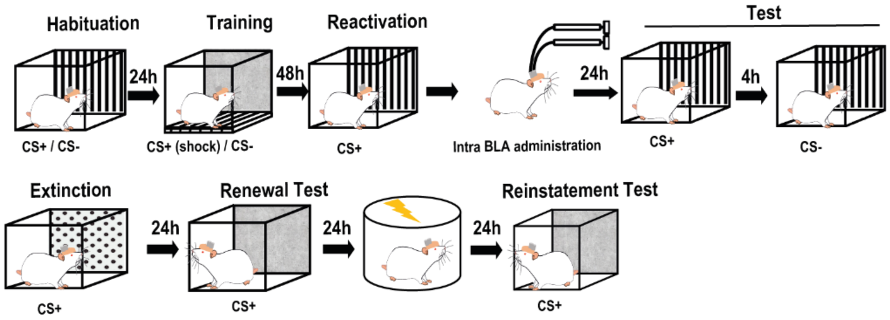


b

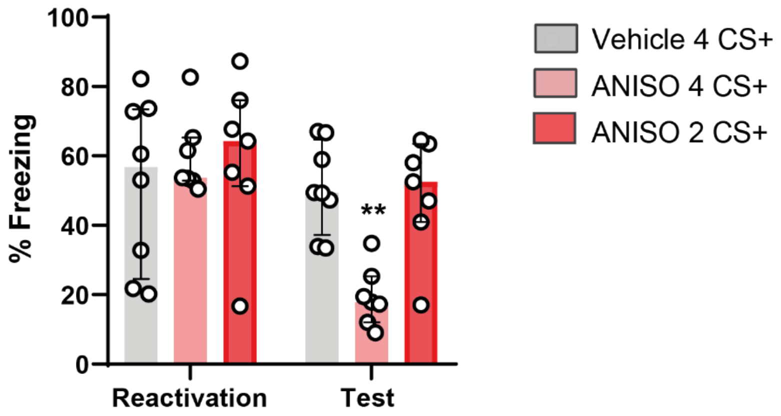


**Supplementary Fig. 2. Standardizing a memory reactivation protocol for inducing reconsolidation.** Only animals exposed to 4 CS+ sessions and intra-BLA anisomycin injection after reactivation exhibited memory impairment during the test (mixed ANOVA: $F (2,19) = 12.82$, $P < 0.001$, group x time interaction) followed by Bonferroni's multiple comparisons post hoc test (Vehicle 4 CS+ vs. ANISO 4 CS+, P = < 0.01; ANISO 4 CS+ vs. ANISO 2 CS+, P = < 0.01).

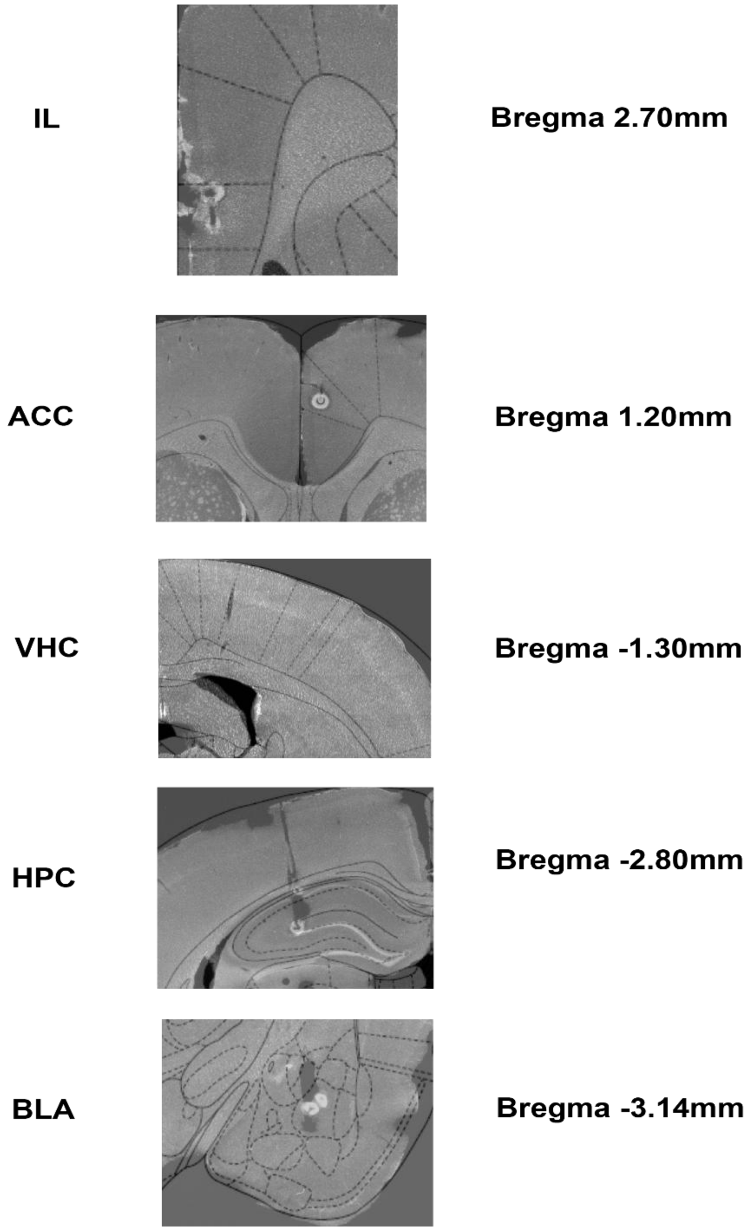


**Supplementary Fig. 3. Histological verification of electrode placements.** A coronal section of the anterior cingulate cortex (ACC), infralimbic cortex (IL) ventral hippocampal commissure (VHC), dorsal hippocampus (HPC) and basolateral amygdala (BLA) stained with DAPI is shown. recording sites on each shank were lesioned (white dots) after the end of the experiment by applying 100 µA anodal direct current for 10 s via electrode tips. The position of the electrodes was maintained throughout all experiments requiring neuronal recording or stimulation.
